# The Five-Primer Challenge: An inquiry-based laboratory module for synthetic biology

**DOI:** 10.1101/745794

**Authors:** Suzie Hsu, Jay Huenemann, Vishwesh Kulkarni, Rodrigo Ledesma-Amaro, Robert Moseley, Nikita Mukhitov, Rashmi Tiwari, Shue Wang, Michael J. Smanski

## Abstract

New technologies in DNA synthesis and assembly give genetic engineers complete freedom in genetic design, where virtually any plasmid DNA sequence can be created efficiently and economically. Learning how to design, construct, and test new DNA sequences is a critical skill for researchers in molecular biology and biotechnology. Here we present a student-centered, inquiry-based module in which students learn how to control bacterial gene expression by appplying various DNA assembly techniques. The central activity in this learning module is termed the ‘Five-Primer Challenge’. Each student is allowed to order up to five 60-mer oligonucleotide primers to then modify a GFP expression plasmid with the goal of increasing GFP expression as much as possible. This module was developed and implemented at the 2016 Cold Spring Harbor Laboratory Synthetic Biology Course, and was effective at engaging students in critical thinking and in promoting student learning.

## Introduction

Inquiry-based learning is a student-focused, active-learning method in which students are confronted with a problem and must design, perform, and analyze experiments to form their own conclusions. The role of an instructor in an inquiry-based learning environment is to facilitate learning and provide information where needed (Savery 2015). Providing students with research-like experiences has been shown to yield superior learning outcomes than traditional teaching methods (Luckie *et al.* 2004), as long as the instructor carefully adapts activities to account for the students’ incoming knowledge and skill level (Kirschner *et al.* 2006). Wet-lab experiments and inquiry-based projects can reinforce material learned during didactic lectures and provide a venue for practicing the formation of hypotheses, design of experiments, and critical analysis of data (Myers and Burgess 2003). Learning modules that incorporate several styles of instruction accommodate students with diverse learning styles and promote teamwork skills such as brainstorming, scientific discussion, and decision-making, which are difficult to teach solely via lecture-based instruction. Engaging students actively in problem-solving and experimentation is an important component of science education. Inquiry-based laboratory curricula have been successfully applied to teach molecular biology (Bugarcic *et al.* 2012) and biochemistry (Gray *et al.* 2015) in the past, but additional modules are needed to address rapid technological progress in these areas and the emerging field of synthetic biology.

A prominent example of inquiry-based learning in the synthetic biology community is the BioBricks foundation and the international Genetically Engineered Machines (iGEM) competition (Smolke 2009; Mitchell *et al.* 2011). The objective-oriented team competition gives students first-hand experience in hypothesis-driven research, and the foundation provides access to genetic reagents to make this experience economically viable for hundreds of schools around the world, and students ranging from high school to graduate school participate(Mitchell *et al.* 2011). However, the iGEM competitions require significant time and resource investment to mentor students, oversee research, develop project presentations, and travel to competitions. Also, the number of participants in iGEM teams at most universities is a small fraction of the population majoring in biotechnology related fields. Inquiry-based lab experiments that are more easily incorporated into core lab-course curricula are needed to provide the majority of students with similar inquiry-driven research experience. Ideally, these modules would retain aspects of student-driven hypothesis creation and experimental design as they are scaled-up to accommodate more students.

DNA synthesis and assembly methods have rapidly advanced in the past decade and represent a valuable skill set for new students trained for careers in biotechnology (Ellis *et al.* 2011). With low cost and rapid turnaround time for DNA synthesis, it is feasible to incorporate student-driven genetic engineering experiments in an educational setting. In one notable example, an undergraduate course was taught *de novo* synthesis of DNA in a semester-long inquiry-based setting (Dymond *et al.* 2009). We sought to build on this effort by developing a learning module where students apply knowledge gained through didactic lectures on the control of gene expression to design experiments that require DNA synthesis and assembly towards a realistic genetic engineering goal.

In the following sections, we describe a novel, inquiry-based learning module to teach students fundamental aspects of engineering bacterial gene expression and introduce them to several practical DNA assembly methods: Isothermal (Gibson) Assembly, Golden Gate Assembly, and PCR ligation. The immediate learning objectives of the module focus on understanding the principles and concepts of each DNA assembly method and learning protocols for setting up reactions as well as analyzing experimental results. At a higher level, students learn the advantages, limitations, and failure modes of each method so that they can apply them to support large-scale DNA assembly projects. Ultimately, the knowledge laid the foundation for the students to create and validate DNA assembly solutions to the Five-Primer Challenge. This module was conceived and demonstrated at the Cold Spring Harbor Laboratory Summer Course in Synthetic Biology (https://cshlsynbio.wordpress.com/). The aim of this publication is to provide an architecture for a fun, engaging, inquiry-based teaching module that can be taught either as part of a rigorous one-week short course, or used as a component in a semester-long laboratory course in synthetic biology or genetics.

## Materials and Methods

### Project Participants

The Cold Spring Harbor Synthetic Biology Course is a two-week long course, taught yearly since its founding in 2013, that focuses on exposing students to state-of-the-art synthetic biology techniques and methods. The course is broken up into two sections; the first week consists of one-day modules that entail a balance of lecture and lab work on various synthetic biology topics (*e.g.* DNA assembly methods, biological modeling, *in vitro* transcription/translation systems, etc), while the second week involves the students participating in more intensive laboratory-based modules. The module described here was designed to be implemented in a rigorous short-course, however the content could easily be delivered in a less intensive course, spread out over many weeks.

The sixteen students in the 2016 course were at various stages in their careers and included PhD students, industry professionals, and academic assistant professors. Skill levels ranged from no previous wet-lab experience to multiple years of experience in genetic engineering. The low student:teacher ratio (4:1) for a particular module allowed for individual attention to be given to students at differing ability levels.

### Learning objectives

The measurable learning objectives for the module as a whole are as follows:

1. Students are able to design genetic constructs at a high level of abstraction (*e.g.* plasmid map) that encode a bacterial expression unit, taking into consideration *cis*-regulatory elements required for efficient transcription and translation.
2. Students are able to design, at the DNA sequence level, rational mutations to a bacterial expression unit towards an objective function of increasing or decreasing transcription and/or translation rates of the encoded gene.
3. Students are able to design oligonucleotide primers to be used in a Gibson DNA assembly reaction that will allow (i) the seamless fusion of two distinct genetic elements, or (ii) the introduction of designed mutations at or near the junctions of two distinct genetic elements.
4. Students are able to design oligonucleotide primers to be used in a Type IIS (Golden Gate assembly reaction that will allow (i) the seamless fusion of two distinct genetic elements, or (ii) the introduction of designed mutations at or near the junctions of two distinct genetic elements.
5. Students are able to design oligonucleotide primers to be used in a PCR-ligation reaction that will produce allow the introduction of designed mutations at or near the ligation junction.
6. Students can design a DNA assembly plan to create a plasmid of interest, taking into consideration the strengths, weaknesses, and failure modes of Gibson, Golden Gate, and PCR-ligation reactions to select appropriate methods.

### Module design

At the outset of the module, students were introduced to the Five-Primer Challenge. Foundational knowledge and wet-bench skills required to complete the challenge were established through a series of white-board lectures, mini-labs, and literature discussions (Table 1). During the Cold Spring Harbor Short Course, mini-lectures and labs took place during the first two days of the module (typically a different lecture/lab combination each morning and afternoon section). On the third day, students were given the morning to design their DNA assembly strategies and order their five oligonucleotides, and the afternoon was spent in a literature discussion/lecture focused on integrating the molecular methods to create an overall DNA assembly pipeline tailored to a specific project. Students used the fourth day to build their plasmids and transform an *E. coli* expression host, and all recombinant strains were cultured and analyzed on the fifth day by fluorescent plate reader or flow cytometry.

**Table 1.**
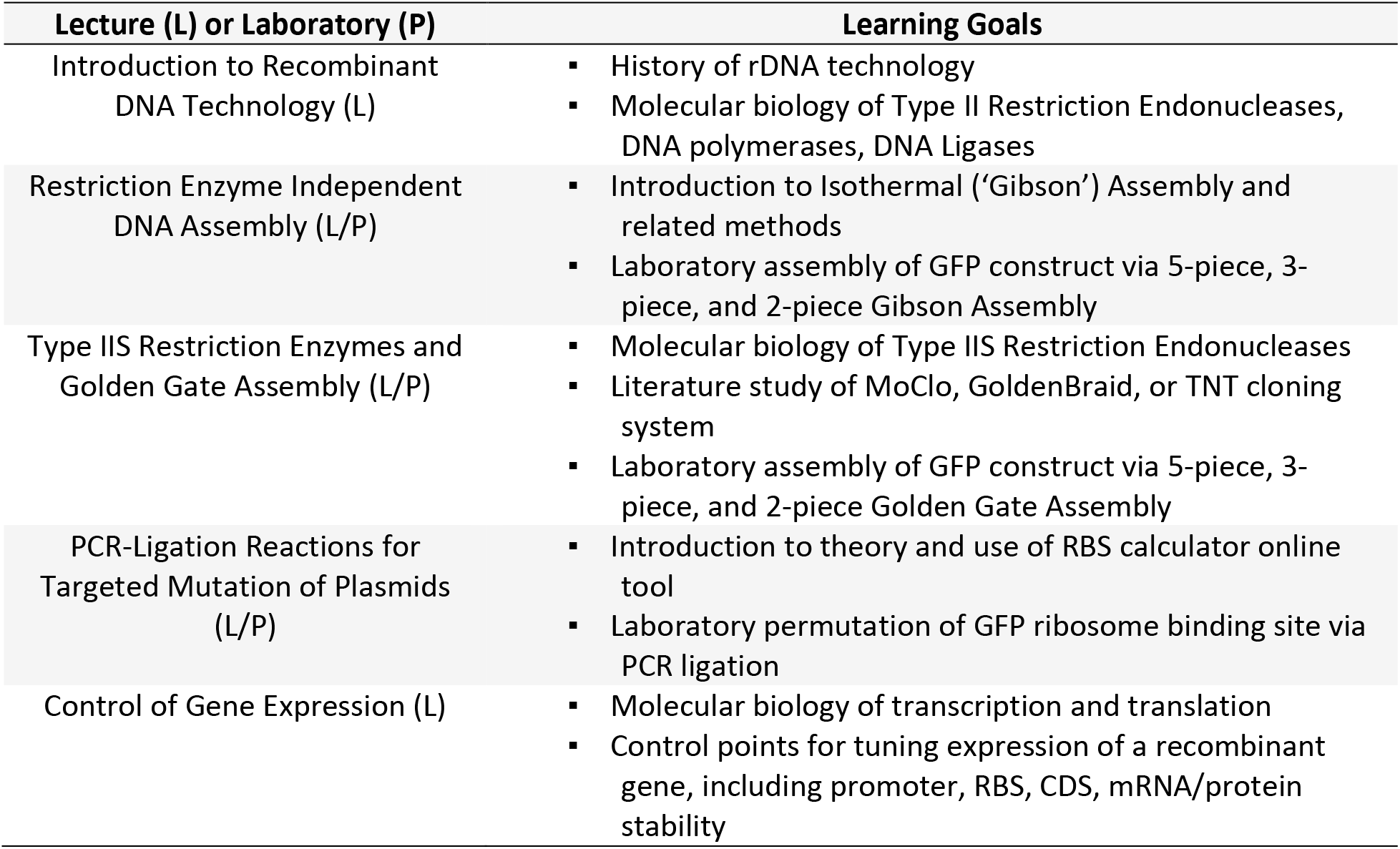
Lecture and lab topics and corresponding learning goals(Handelsman *et al.*2007)

| Lecture (L) or Laboratory (P) | Learning Goals |
| --- | --- |
| Introduction to Recombinant DNA Technology (L) | History of rDNA technology<br>Molecular biology of Type II Restriction Endonucleases, DNA polymerases, DNA Ligases |
| Restriction Enzyme Independent DNA Assembly (L/P) | Introduction to Isothermal ('Gibson') Assembly and related methods<br>Laboratory assembly of GFP construct via 5-piece, 3-piece, and 2-piece Gibson Assembly |
| Type IIS Restriction Enzymes and Golden Gate Assembly (L/P) | Molecular biology of Type IIS Restriction Endonucleases<br>Literature study of MoClo, GoldenBraid, or TNT cloning system<br>Laboratory assembly of GFP construct via 5-piece, 3-piece, and 2-piece Golden Gate Assembly |
| PCR-Ligation Reactions for Targeted Mutation of Plasmids (L/P) | Introduction to theory and use of RBS calculator online tool<br>Laboratory permutation of GFP ribosome binding site via PCR ligation |
| Control of Gene Expression (L) | Molecular biology of transcription and translation<br>Control points for tuning expression of a recombinant gene, including promoter, RBS, CDS, mRNA/protein stability |

### Assessment of Student Learning

Student learning was measured using a combination of informal and formal techniques. Content-driven white-board lectures were interrupted with multiple-choice questions (Supplementary Information) to gauge students’ understanding of key concepts and engage students in critical thinking. Homework assignments were practically oriented and included (i) the manipulation of DNA sequence files using plasmid editing software (ApE, http://biologylabs.utah.edu/jorgensen/wayned/ape/) (Assignment 1, Supplementary Information), (ii) the design of oligonucleotide primer sequences that could be used to achieve a desired DNA assembly product (Assignment 1, Supplementary Information), and (iii) use of the RBS Calculator (https://www.denovodna.com/software/) to design ribosome binding sites of various strength (Assignment 2, Supplementary Information). Misconceptions were addressed immediately with students in small group settings. Critical thinking and problem solving capabilities were informally assessed by reviewing student-proposed solutions to the Five-Primer Challenge. Successful solutions were judged by (i) the likelihood that the plasmid design changes would reasonably lead to increased cellular fluorescence, and (ii) whether the oligonucleotide sequences and proposed DNA assembly strategy were feasible to achieve the plasmid design changes. Rank-order results from the Five-Primer Challenge did not factor into assignment of grades, but a symbolic prize was given to the winner. A post-course survey was administered to gather student feedback on course and learning objectives, but no formal final examinations were given during the CSHL Synthetic Biology Course.

### Experimental methods

Detailed description of the experimental materials and methods used for laboratory work described in this manuscript can be found in the Supplementary Materials.

## Results

### Demonstration of student learning in primer design activities

The student’s ability to successfully design primers for a Gibson Assembly was assessed before and after participation in the Five-Primer Challenge. Immediately following the mini-lecture on Gibson Assembly, students from Session 2 were supplied with electronic files containing annotated sequence files for two plasmids: one was a GFP-expression plasmid containing a kanamycin resistance marker, and a second containing an ampicillin resistance marker. Students were asked to generate four oligonucleotide primers that could be used to amplify DNA fragments from these plasmids that could be assembled via a Gibson Reaction to replace the kanamycin marker with an ampicillin marker in the opposite orientation (*i.e.* transcribed from the opposite strand as the kanamycin gene, but in the same position on the plasmid). Three of the five students produced primer designs that would not produce dsDNA fragments that would properly assemble in a Gibson reaction. Two of the five students produced primer designs that would properly assemble, but would not lead to functional expression of the new resistance cassette, as the cis-acting regulatory elements were positioned inappropriately. These incorrectly designed primers were used as a teaching tool in a follow-up lecture, and the reasons why they would fail were discussed in detail. At the end of the 4-day teaching module, students were given a similar problem and 5/5 produced appropriate primer designs.

### Comparing DNA assembly efficiencies in a model system

Students performed two lab exercises on day one, comparing Golden Gate and Gibson Assembly reactions performed with 2, 3, or 5 DNA fragments. Each assembly reaction produced the same GFP expression plasmid, which would only result in GFP production in correctly assembled plasmids. Because of this, *E. coli* colonies could be screened visually using a transilluminator to determine reaction efficiency (Figure 2), and costly validation by plasmid sequencing could be avoided. For both Golden Gate and Gibson Assembly reactions, two- and three-piece reactions were more efficient than five-piece reactions, with Golden Gate performing slightly better than Gibson for the more complex assemblies (Figure 2). During the CSHL short course, a parallel experiment aiming to minimized reaction volumes was performed where total reaction volumes ranged from 2 μL to 125 nL. Precision mixing of substrate fragments and reagents was achieved with a LabCyte Echo liquid handler. Successful reactions were seen for each of two-, three-, and five-piece assemblies for total reaction volumes as low as 250 μL, with similar efficiencies as seen for the manually pipetted, 10 μL reactions (data not shown).

**Figure 1.**
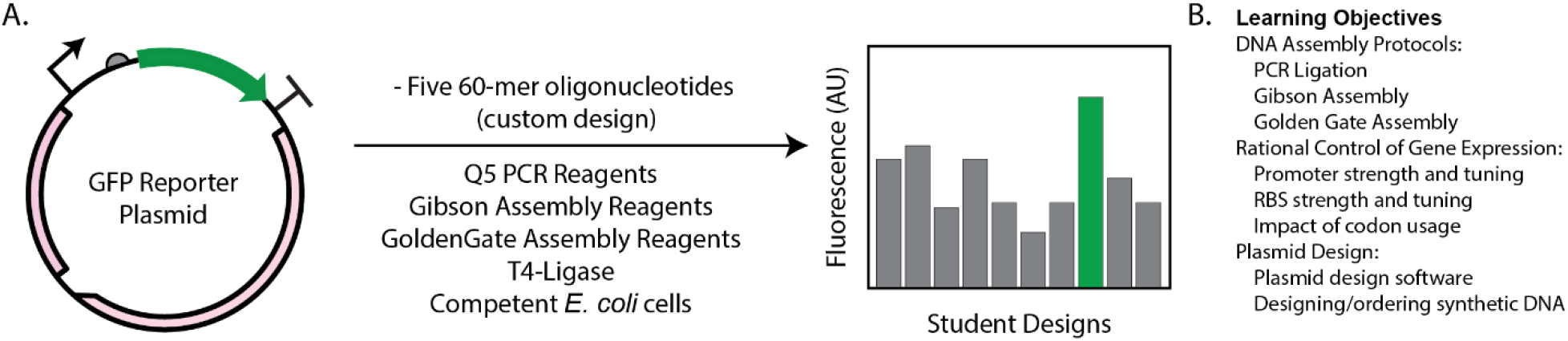
Summary of Five-Primer Challenge and learning objectives. (A) Reagents that are available to students are listed, and sample graph comparing re-designed reporter plasmids is shown at right. (B) Learning objectives are listed by category.

**Figure 2.**
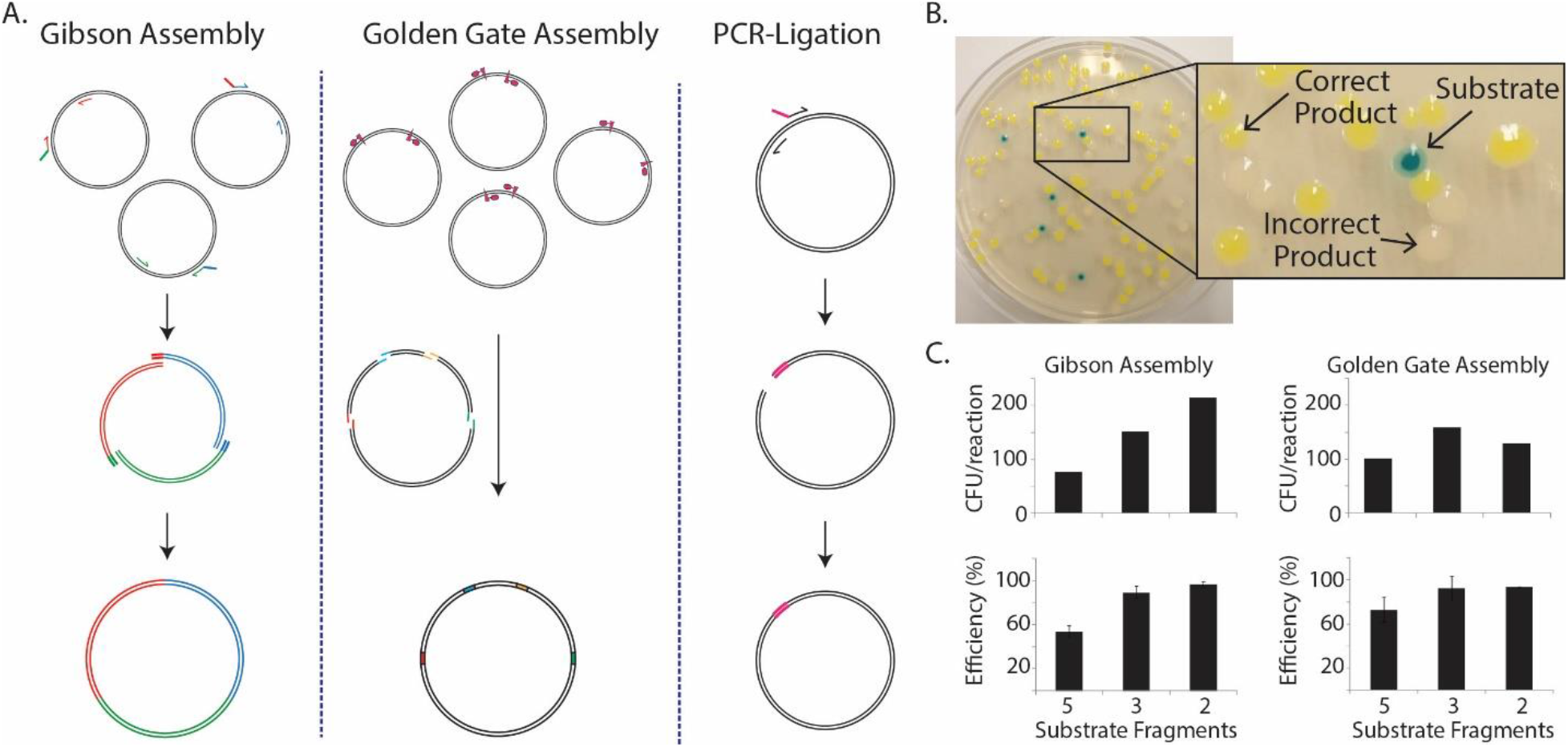
Sample data from practical lab experiments. (A) Schematic illustration of three laboratory protocols taught during lab/practical component of module. (B) Scoring correctly assembled (green) vs incorrectly assembled (white) product plasmids using a visual screen. In practice, colonies are screened at a much smaller size using a UV transilluminator to more easily discriminate fluorescent vs. white colonies. (C) Quantitative comparison of reaction efficiencies for Gibson (left) and Golden Gate (right) assembly reactions. Efficiency is defined as the GFP-expressing colonies divided by the number of total colonies. Error bars for reaction efficiency denote one standard deviation from the mean, from five independent replicates.

### Students learn and apply diverse strategies to control protein production

The nine students who participated in the Five-Primer Challenge (four in session 1, five in session 2), were given complete freedom of genetic design to engineer a highly fluorescent strain of *E. coli*. The diverse solutions they designed serve as evidence to their engagement in creative problem solving. The solutions, summarized in Figure 3 and Table 2, included manipulation of promoter strength, ribosome binding site strength, gene copy number, and synonymous mutations in the GFP CDS to reduce secondary structure during translation initiation. Many solutions used tools such as the Ribosome Binding Site Calculator that were introduced during the mini-lectures. However, a number of the students performed substantial literature review to identify additional control points that could give them an edge in the Challenge. These included non-synonymous point mutations in the GFP CDS to increase the fluorescence per protein molecule, and mutations to the plasmid origin of replication that would lead to ‘runaway replication’ and dramatically increase the copy number of the expression construct. A third category consisted of ideas that were not directly based on literature precedent, but were sound and worthy of experimental validation (Table 2). In surveys collected by Cold Spring Harbor Laboratories following the course, several students noted that the ‘thought experiments’ that were required to design solutions to the Five-Primer Challenge forced them to engage with the material more deeply than in previous molecular biology courses.

**Figure 3.**
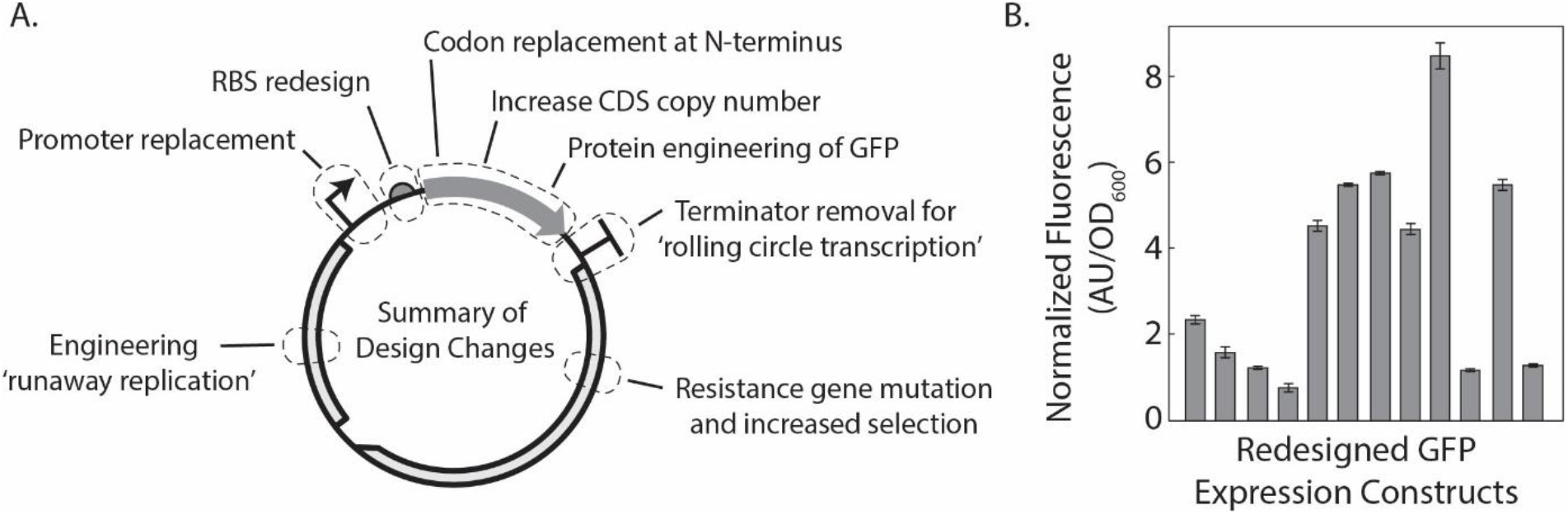
Five-Primer Challenge results. (A) Summary of student solutions to increasing expression of recombinant GFP in *E. coli*. (B) Plate-reader results of normalized fluorescence for student and instructor designed expression constructs. Bars represent mean fluorescence of two technical replicates with error bars showing range of fluorescence readings. Expression results are not connected to design changes for purpose of future competitions.

**Table 2.** Student solutions to the Five-Primer Challenge

| Discussed in mini-lectures | Ideas generated based on literature review | Ideas generated without literature precedent |
| --- | --- | --- |
| Changing GFP promoter<br>Re-designing GFP ribosome binding site.<br>Synonymous mutations in N-terminal region of GFP CDS.<br>Changing relative order/orientation of features in expression plasmid. | Non-synonymous mutations in GFP that increase fluorescence per molecule.<br>Mutations in origin of replication that lead to 'runaway replication'.<br>Addition of UP element upstream of -35 of promoter region | Removing all transcriptional terminators to yield long, poly-GFP mRNA from circular plasmid.<br>Decrease strength of resistance marker and grow on high antibiotic concentration to select for rare mutants with strong expression from plasmid.<br>Increase copy number of GFP to arbitrarily long constructs with creative DNA assembly protocol. |

### Assessment of student Five-Primer solutions

Students each selected an *E. coli* transformant for growth and fluorescence analysis in a 96-well plate-reader. Technical replicate measurements of GFP fluorescence were made for each transformant and are reported in Figure 3B. Student GFP reporter constructs varied in expression strength over one order of magnitude. Time limitations at the CSHL short course prevented biological replicates to be measured over multiple days, but this should be incorporated to future iterations of the Five-Primer Challenge. Results in Figure 3B are purposely not linked to genetic designs in order to encourage experimentation and creativity in the future.

## Discussion

Improvements in cost, speed, efficiency, and flexibility of DNA synthesis and assembly methods now provide unprecedented freedom for designing and fabricating genetic constructs. Genetic engineers are no longer constrained by historical precedent to use one of a small number of existing expression plasmids to overproduce a protein of interest. Instead, the ability to order synthetic DNA and have it delivered next-day allows genetic engineers to easily change any aspect of a construct’s design.

The next generation of genetic engineers need a strong foundation in both DNA synthesis/assembly methods as well as a mechanistic and theoretical understanding of gene expression; they need to know what to design and how to build it. To address this pedagogical need, we have developed and demonstrated a fun, engaging, and defined teaching module that fosters student learning and creative problem solving. Named the ‘Five-Primer Challenge’, this teaching module provides students with the knowledge-base and resources to rationally design, build, and test a GFP expression plasmid, with the aim of increasing total cellular fluorescence. Decreasing costs of DNA synthesis (the five primers cost ~$50 per student for custom synthesis) enabled this experiment to be run as a competition, with each student responsible for using oligonucleotides of their own design to produce a re-engineered plasmid.

The merit of classroom competitions is debated in the pedagogical literature (Cantador and Conde 2010). Supporters of classroom competitions note that they can increase student motivation and learning (Verhoeff 1997), and result in students spending extra effort in learning the material compared to non-competitive environments (Fasli and Michalakopoulos 2005). However, others have argued that classroom competitions generate additional stress on students that outweighs the potential benefits (Vockell 2004). There is general consensus that competitions can be organized in a way to retain the benefits while minimizing the negative aspects, for example competing as small teams, or competing as a class against previous and/or future classes (Yu *et al.* 2002).

Competitions are most common in computer science classrooms where students write custom programs to accomplish a narrowly-defined objective function. In the Five-Primer Challenge, the objective function is for students to create a GFP expression plasmid with the strongest possible expressed levels using the resources available to them. While critics of classroom competitions note that they can place additional stress on student performance that detracts from educational goals, this was not reported by students in the post-class survey. Two possible explanations for this include, first, that formal student assessment/evaluation is not part of the CSHL Synthetic Biology course, so performance in the competition was for ‘bragging rights’ only. If this module is included as part of a curriculum in molecular biology/genetic engineering, we recommend that grades are given based on the plausibility that a student’s design will increase fluorescence, but not based on results of the competition. Second, a team-dynamic was introduced by the instructor claiming that he could win the challenge with only two primers. This created an extra level of competition (all students against teacher) that encouraged students to work together, brainstorming diverse strategies to win the challenge such that at least one would beat the instructor. Because of this, the Five-Primer Challenge had aspects of both a team challenge and an individual challenge.

Proponents of classroom competitions note improvements in time and effort spent as well as motivation and engagement (Fasli and Michalakopoulos 2005). During this module, the students spent time and effort well beyond what was expected, both at night and during meals, to brainstorm and discuss possible solutions to problem. The alternative solutions proposed (Table 2 and Figure 3) are evidence that students understand the major and minor control points for engineering heterologous protein production. Less than half of the solutions proposed were covered during the didactic lectures, with the rest coming from literature surveys or creative thinking (Table 2).

Several aspects of this module design could be examined further in future studies. For example, what is the impact of competition organization (all vs all, class vs instructor, no competition) on (i) the diversity of solutions proposed and (ii) the amount of teamwork displayed by the students? How does participation in the didactic lectures and practical experimental labs affect the quality and diversity of student hypotheses?

The teaching module presented here was effective at motivating and engaging students during the CSHL Synthetic Biology course and can be incorporated into existing laboratory courses aimed at teaching students the fundamentals of molecular biology with an emphasis on biotechnology applications.

## Acknowledgement

We would like to thank Schmidt-Dannert laboratory for generously giving us the pUCBB-NTH6-eGFP plasmid for the DNA assembly module. We would like to thank co-instructors at the CSHL SynBio Course, Vincent Noireaux, Chase Beisel, Mary Dunlop, Mo Khalil, and Harris Wang for help in organizing the course. Lastly we would like to thank CSHL staff members Rachel Lopez, Barbara Zane, and Breanna Demestichas for help in administering the course.

